# Disentangling key species interactions in diverse and heterogeneous communities: A Bayesian sparse modeling approach

**DOI:** 10.1101/2021.07.23.453227

**Authors:** Christopher P. Weiss-Lehman, Chhaya M. Werner, Catherine H. Bowler, Lauren M. Hallett, Margaret M. Mayfield, Oscar Godoy, Lina Aoyama, György Barabás, Chengjin Chu, Emma Ladouceur, Loralee Larios, Lauren G. Shoemaker

## Abstract

Modeling species interactions in diverse communities traditionally requires a prohibitively large number of species-interaction coefficients, especially when considering environmental dependence of parameters. We implemented Bayesian variable selection via sparsity-inducing priors on non-linear species abundance models to determine which species-interactions should be retained and which can be represented as an average heterospecific interaction term, reducing the number of model parameters. We evaluated model performance using simulated communities, computing out-of-sample predictive accuracy and parameter recovery across different input sample sizes. We applied our method to a diverse empirical community, allowing us to disentangle the direct role of environmental gradients on species’ intrinsic growth rates from indirect effects via competitive interactions. We also identified a few neighboring species from the diverse community that had non-generic interactions with our focal species. This sparse modeling approach facilitates exploration of species-interactions in diverse communities while maintaining a manageable number of parameters.

## 2 Introduction

Understanding what maintains the diversity of life—where and how species abundances change through time—has long fascinated and challenged ecologists. It is widely accepted that community composition in any given time and place is driven by the interplay of species interactions, responses to environmental conditions, and feedbacks between local and regional dynamics (Chesson, 2000; HilleRisLambers *et al*., 2012; Vellend, 2020). However, given the myriad of biotic interactions that may, themselves, be mediated by underlying environmental conditions (Bulleri *et al*., 2016; Germain *et al*., 2018; Letten *et al*., 2018), feasibility and model overfitting concerns quickly arise when trying to incorporate observed levels of diversity. Arguably, the magnitude of this methodological limitation has even shaped our historical theoretical frameworks and empirical tests. For example, classic species trait trade-offs, such as the competition-colonization trade-off, apply for species pairs (Levins & Culver, 1971; Tilman, 1982). Similarly, while modern coexistence theory (Chesson, 2000) can be applied to any level of species richness (Spaak & De Laender, 2020), the vast majority of empirical studies focus on pairwise species comparisons (e.g. Kraft *et al*. 2015; Wainwright *et al*. 2019) and the effect of environmental variation on these comparisons (Bimler *et al*., 2018; Lanuza *et al*., 2018). Yet nonlinearity, higher-order interactions, and intransitivity in diverse systems may yield complex dynamics that dramatically alter population growth and coexistence dynamics (Allesina & Levine, 2011; Li *et al*., 2021; May & Leonard, 1975; Mayfield & Stouffer, 2017). The further development and empirical testing of these theories thus requires a statistical approach that is applicable in diverse communities and is capable of identifying and incorporating key species interactions and environmental covariates.

To date, empirical studies of population dynamics and species coexistence frequently take one of two approaches for dealing with parameterization limitations that arise in diverse communities and varied environments. In the first approach, experimental studies focus on a few focal species. For example, Wainwright et al. (2019) examined coexistence based on pairwise interaction coefficients between four annual forbs in two locations and across two water availability treatments—a lofty number of species interaction coefficients to estimate—but still a relatively small subset of the community’s full diversity (10 − 14 species in 0.09 m^2^; Dwyer *et al*. 2015). Finer-scale environmental variation can further limit the number of species that can be feasibly incorporated: in a study of grass and forb coexistence under variable rainfall regimes, Hallett et al. (2019) considered four rainfall conditions, requiring estimates of eight distinct species interaction coefficients even with only two species. Isolating species interactions across environmental conditions is a high barrier in species rich communities, even in laboratory and microcosm based studies (Letten *et al*., 2018).

In the second approach, often used to interpret observational data, species are grouped into broad categories. At the most extreme, a single interaction coefficient is then calculated between the focal species and all heterospecific individuals—regardless of their identity (Clark *et al*., 2020a; Uriarte *et al*., 2004). Heterospecifics may also be grouped more finely, for example, according to their taxonomic relationship (Uriarte *et al*., 2004) or their origin status and life form (e.g. native versus exotic and grasses versus forbs) (Martyn *et al*., 2020). Alternatively, functional groups can be created by grouping species according to their traits (e.g. specific leaf area, canopy height, seed number) (Kühner & Kleyer, 2008; Uriarte *et al*., 2004). However, this methodological approach often necessitates *a priori* knowledge of the system and makes an underlying assumption that species grouped together will interact similarly with each other and with the focal species. These assumptions are often not met (Mayfield & Levine, 2010), suggesting a need for a more parsimonious and robust methodology that would allow the data to inform species groupings.

Various alternative statistical approaches have been proposed to assess species interactions using observational data. For example, joint species distribution modeling has become a common approach to infer species interactions from co-occurrence patterns (Legendre & Gauthier, 2014; Ovaskainen *et al*., 2019, 2017b). However, in addition to species interactions, patterns of co-occurrence may result from environmental sorting (Barner *et al*., 2018), or dispersal patterns (Schamp *et al*., 2015). Further, co-occurrence patterns are scale dependent and regional analyses are not suited to assessing local-scale species interactions (König *et al*., 2021). Recognizing a need to directly estimate species interaction coefficients, recent work has expanded multivariate autoregressive models for use in more diverse communities (Picoche & Barraquand, 2020), including examining which linear combinations of species abundances best predict future growth rates (Ovaskainen *et al*., 2017a). This approach is effective for binning species based on their competitive effects, but does not account for variation in the environment. Clark et al. (2020b) recently developed a state-space hierarchical Bayesian model to assess the effect of environmental gradients on nonlinear species abundance patterns, incorporating environment responses in species’ density-independent growth rates, but not in species interactions (Clark *et al*., 2020b). Lastly, García-Callejas et al. (2020) developed a method to incorporate environment responses in species’ density-dependent growth rates but without the flexibility of Bayesian approaches. Independently, these different methodological developments each address one of the largest hurdles for modeling species abundances in diverse communities: (1) identifying important species interactions and (2) accounting for the mediating effect of the environment (here referred to as species-environment interactions). Addressing these two aspects simultaneously would solidify a path forward for characterizing species interactions in diverse communities and across environmental gradients.

Here, we present an approach for modeling dynamics in diverse communities and across environmental gradients. The approach balances realism and complexity without extensive experimental manipulation or *a priori* assumptions regarding species groupings. Our method is based on two innovations to standard population and community ecology models. First, we define heterospecific species interaction coefficients as linear combinations of the average interaction strength and species-specific deviations from this average. In parallel, we allow environmental covariates to modify species intrinsic growth rates and the strength of biotic interactions—both the average and species deviation terms. We implement this approach using a Beverton-Holt model of community dynamics (Beverton & Holt, 1957) within a single growing season, although the method can easily be adapted to other models of population abundance (e.g. Mayfield & Stouffer 2017; Ricker 1954) or incorporate additional dynamics such as seed banks or dispersal (Levine & HilleRisLambers, 2009; Thompson *et al*., 2020). Second, we extend Bayesian statistical methods for variable selection via sparsity-inducing priors in linear models (such as Lasso and Ridge regression; Hastie *et al*. 2015; Piironen *et al*. 2017) to our non-linear abundance model, thereby reducing the number of terms included in the final model fit, yielding a ‘sparse model.’ By coupling these two modeling approaches, we can identify heterospecific species that deviate in their interaction strength, and how environmental gradients alter species’ density-independent growth rates and biotic interactions. We explore model effectiveness using simulated data and apply the model to empirical data from a highly diverse (45 species) annual plant community.

## 3 Methods

### 3.1 Deconstructing species interaction coefficients and fecundity

Models of community dynamics incorporate species-specific interaction coefficients for each species pair, commonly denoted as *α_i,j_*, the effect of species *j* on species *i*, resulting in a large number of parameters required to model diverse communities or environmental relationships. To reduce the number of parameters required to model diverse communities and incorporate environmental variation, we start with a partitioning approach. We first define these interaction terms as

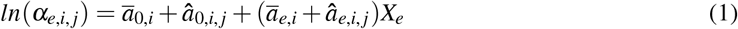

where *α_e,i,j_* is the effect of species *j* on species *i* in environment *e* with *i* ≠ *j*. In Eqn. 1, *ā*_0,*i*_ is the effect of an average heterospecific individual on individuals of species *i*, *â*_0,*i*,*j*_ is the deviation from this average effect associated with species *j*, *ā_e,i_* is the average slope of species *i*’s interaction coefficients with environmental covariate *X_e_*, and *â*_*e*,*i*,*j*_ is the deviation from this slope associated with species *j*. Upon first glance, using Eqn. 1 may seem counter productive as it increases the number of parameters compared to traditional interaction coefficients. However, in the next section we describe how coupling this approach with sparsity inducing priors in a Bayesian context can dramatically reduce the number of required parameters by identifying only the necessary species-specific terms (*â*_0,*i*,*j*_ and *â*_*e*,*i*,*j*_) for accurately modeling population dynamics of species *i*.

While intraspecific competition (*α_e,i,i_*) could in principle be modeled according to Eqn. 1, we instead define it separately as:

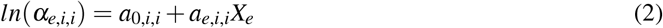

where *a*_0,*i*,*i*_ and *a_e,i,i_* are the intercept and slope for the effect of intraspecific individuals. As both theoretical expectations (Chesson, 2000) and empirical results (Adler *et al*., 2018) point to the importance of intraspecific competition, we use Eqn. 2 to explicitly exclude the intraspecific terms from the sparsity inducing process defined in the next section. These terms, therefore, will always be included in the final model fit.

Interaction coefficients (Eqns. 1 & 2) can be incorporated in many different models of community dynamics. We use the Beverton-Holt model due to its legacy in studies of annual plant communities and coexistence theory (e.g. Godoy & Levine 2014; Kraft *et al*. 2015). We emphasize, however, that our general statistical approach can be adapted to other population models. In the Beverton-Holt model, the fecundity *F_e,i_* of a focal species *i* in environment *e* is modeled as:

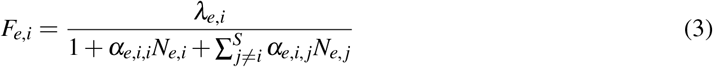

Fecundity depends on a species’ intrinsic growth rate (i.e. density-independent seed production; *λ_e,i_*) and the competitive effects of all *S* species in the community (*α_e,i,j_* terms as defined by Eqns. 1 &2) scaled by each species’ abundance (*N_e, j_*) (Levine & HilleRisLambers, 2009; Pérez-Ramos *et al*., 2019; Shoemaker & Melbourne, 2016). To incorporate environmental variation in intrinsic growth rates, we model *λ_e,i_* as:

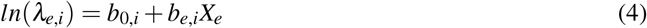

where *b*_0,*i*_ is the intercept of the intrinsic growth rate and *b_e,i_* its slope with environmental covariate *X_e_*. We use Eqn. 3 to model observed fecundity within a single growing season of both simulated and empirical data.

### 3.2 Incorporating sparsity-inducing priors

By deconstructing interaction coefficients into a combination of species-specific and generic terms, we can determine which, if any, species-specific terms are necessary for the final model. Allowing only a subset of parameters to take non-zero values is referred to as ‘sparse modeling,’ and various techniques exist to induce sparsity in linear models (Hastie *et al*., 2015; O’Hara *et al*., 2009).

To extend a sparse modeling approach to our non-linear model of fecundity (Eqn. 3), we employ sparsity-inducing priors which act to shrink all but a subset of parameters to 0, thus producing a sparsely parameterized model. Specifically, we model *â*_0,*i*,*j*_ and *â*_0,*i*,*j*_, the species-specific intercepts and slopes of the inter-specific interaction coefficients (Eqn. 1), with regularized horseshoe priors which more accurately estimate large parameter values compared to other sparsity-inducing priors (Bhadra *et al*., 2019; Carvalho *et al*., 2009; Piironen *et al*., 2017; Van Erp *et al*., 2019). Parameters *â*_*i*,*j*_ and *â*_*e*,*i*,*j*_, are given priors Normal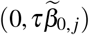 and Normal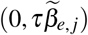 respectively. (Note that since we fit the model for a single focal species, we drop the *i* subscript from the priors for simplicity.) In these priors, *τ* defines the global tendency towards sparsity through its effect on the priors’ standard deviations. In other words, with smaller values of *τ*, the priors for all *â*_*i*,*j*_ and *â*_*e*,*i*,*j*_ parameters become more tightly centered on 0. Conversely, the 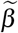 terms allow specific parameters to escape this global trend towards sparsity. As an individual 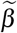 term becomes large, its associated prior becomes wider, and that species-specific term is more likely to be included in the final model. In the regularized horseshoe prior, these 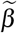 terms are defined as:

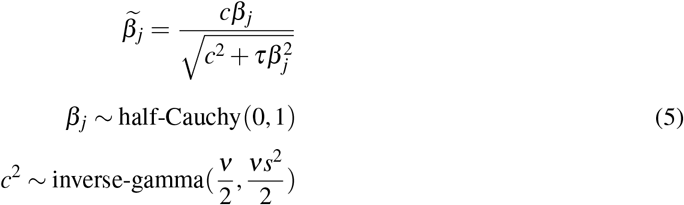

Defining 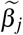 as the combination of a half-Cauchy and inverse-gamma distribution causes large coefficients to be shrunk towards 0 by a Student’s *t* distribution with *ν* degrees of freedom and a scale of *s*^2^ (Piironen *et al*., 2017; Van Erp *et al*., 2019). Following the recommendations of Piironen and Vehtari (2017), we set *ν* to 4 and *s*^2^ to 2. Rather than setting the global shrinkage parameter *τ* to a fixed value, we give it a half-Cauchy prior with scale parameter equal to 1 (*τ* ~ half-Cauchy(0, 1)) and allow the data to inform the posterior distribution of *τ* (Piironen *et al*., 2017; Van Erp *et al*., 2019).

We employ a hybrid approach in which we first fit the full model with regularized horseshoe priors to induce sparsity in the species-specific terms; we subsequently fit a final model using traditional, non-sparse methods. From the preliminary model fit, we identify which species-specific terms have sufficient evidence to be included in the final model fit. We calculate credible intervals (CIs) for each species-specific term in the preliminary model and include in the final model only those terms whose intervals do not overlap 0. By using this approach, we can directly adjust how conservative we wish to be in including model parameters, balancing model prediction, the proportion of variance explained, and simplicity depending on modeling goals (Tredennick *et al*., 2021) (i.e. using a 50% CI will lead to models including more parameters than if we use a 95% CI). Then, for the final model fit, the included species-specific terms (*â*_0,*i*,*j*_ and *â*_*e*,*i*,*j*_) are given standard normal priors (i.e. Normal(0, 1)). In both preliminary and final model fits, the terms defining *λ_e,i_* (*b*_0,*i*_ and *b_e,i_*; Eqn 4) are also given standard normal priors. The intercept and slope terms defining intraspecific competition (*a*_0,*i*,*i*_ and *a_e,i,i_*; Eqn.2) and the generic intercept and slope defining interspecific competition (*ā*_0,*i*_ and *ā_e,i_*; Eqn. 1) are both given weakly informative priors in each model fit, matching the expected scale of these interaction coefficients: *a*_0,*i*,*i*_ ~ Normal(−6, 3), *ā*_0,*i*_ ~ Normal(−6, 3), *a_e,i,i_* ~ Normal(0, 0.5), and *ā_e,i_* ~ Normal(0, 0.5). All models were fit using the stan language with the rstan package (version 2.18.2; Stan Development Team 2018 in R (version 3.5.3; R Core Team 2019). All code for the analyses and simulations presented here can be found at https://github.com/tpweiss06/SparseInteractions.

### 3.3 Simulation tests of model performance

To test our ability to predict changes in population size and recover true parameter values, we first paired our Bayesian sparse modeling approach with simulated Beverton-Holt data using Eqns. 1–4. For the simulations, we generated communities of 15 species in different plots, where each plot was a unique run of the simulation for a given community with a given environmental condition *X_e_*. We aimed to generate population growth rates comparable to those found in a community adapted to its environment. Each species was assigned an intrinsic growth rate *λ_e,i_* following Eqn 4 (Table 1), pairwise species competitive interactions *α_e,i,j_* were composed of the generic competition term *ā*_0,*i*_ with small amounts of variation and a generic environmental response *ā_e,i_*. Seven randomly selected species also had a non-generic competition term through species-specific deviations from *ā*_0,*i*_ (*â*_0,*i*,*j*_). Seven separately selected species had a non-generic environmental response through species-specific deviations *â*_*e*,*i*,*j*_. Intraspecific competition *ā*_0,*i*,*i*_ was set as a fixed value higher than interspecific competition to minimize extinction in the simulations (Table 1). Each plot simulation was run deterministically for 20 time steps with each time step *N*_*t*+1_ = *F_t_N_t_* using *F_t_* from equation 3. This resulted in some subset of the 15 species remaining with populations greater than zero in each plot. Then each population was perturbed by drawing from a normal distribution with mean and standard deviation equal to the previous population size, truncated at 0 to prevent negative population sizes. This perturbed state and the following time step generated our simulated ‘full-community’ data. In addition to 500 full-community plots, we simulated 500 ‘no-competition’ treatments with a single phytometer individual of the focal species per plot, running the Beverton-Holt function for one time step. This simulated treatment matches methods commonly used in experimental studies to parse intrinsic growth rates from competition parameters (Hallett *et al*., 2019; Wainwright *et al*., 2019). Simulation details are included in Supplement 1.

**Table 1:** Parameter components and distributions used in simulations

| Parameter |  | Distribution | Unique to |
| --- | --- | --- | --- |
| $S$ | regional species pool | 15 species | single value for community |
| $N_{i,0}$ | initial population | Gaussian (mean 80, sd 50) truncated at 0 | species, plot |
| $X_e$ | plot environment | Gaussian (mean 0, sd 1) | plot |
| $\lambda_{e,i}$ | intrinsic growth with monotonic environmental response | $e^{b_{0,i}+b_{e,i}X_e}$ | species |
| $b_{0,i}$ | environmentally-independent growth rate | uniform (0 to 1.5) | species |
| $b_{e,i}$ | environmental response in $\lambda$ | Gaussian (mean 0, sd 0.5) | species, plot |
| $\alpha_{e,i,j}$ | competitive interactions with environmental effects | $e^{\bar{a}_{0,i}+\hat{a}_{0,i,j}+(\bar{a}_{e,i}+\hat{a}_{e,i,j})X_e}$ | species pair, plot |
| $\bar{a}_{0,i}$ | generic interspecific competition | Gaussian (mean -7, sd 0.1) | species pair |
| $\hat{a}_{0,i,j}$ | deviation in competition | uniform $\pm(1.5$ to $3.2)$ | 7 species $j$ |
| $\bar{a}_{0,i,i}$ | intraspecific competition | -3.5 | constant |
| $\bar{a}_{e,i}$ | generic environmental variation in competition | Gaussian (mean 0, sd 0.3) | single value for community |
| $\hat{a}_{e,i,j}$ | specific environmental variation in competition | uniform $\pm(0.5$ to $1.5)$ | 7 species $j$ |

We used these simulations to measure our sparse modeling approach’s ability to predict population growth in diverse communities and recover underlying parameters. We selected one focal species and tested our model’s performance using varying numbers of full-community and no-competition plots. We tested out-of-sample predictions on 200 full-community plots not used to fit the model. We then calculated the posterior distribution of the root-mean-square error (RMSE) of model predictions compared to true values for each model fit. This allowed us to quantify the gain in predictive accuracy resulting from including more data.

### 3.4 Empirical application

We additionally applied our model to species interactions and their environmental dependencies in the annual plant understory of the York gum (*Eucalyptus loxophleba* Benth) - jam (*Acacia acuminata*) woodlands of southwestern Western Australia. This community is highly diverse and heterogeneous, with local composition of annual forbs and grasses influenced by gradients in soil nutrients and shade from York gum and jam trees (Dwyer *et al*., 2015; Lai *et al*., 2015). We focused on two York gum-jam woodland remnants: West Perenjori Nature Reserve (29°47’S, 116°20’E) and Bendering Nature Reserve (32°23’S, 118°22’E). Both sites experience a Mediterranean climate with mild winters and long, dry summers (Suppiah *et al*., 2007) and have high overlap in annual species composition, sharing several dominant species. Data used for this study were originally collected as part of a larger experiment described in full in Wainwright et al. (2019). We focus on two species used as focal species in the original study and common to both reserves: *Waitzia acuminata*, an abundant native annual forb, and *Arctotheca calendula*, a prevalent exotic annual forb.

We used data from 11 experimental blocks in Bendering Nature Reserve and 18 blocks in West Perenjori Nature Reserve. Each block was ≈ 15 × 15 m, a size selected to account for previously identified soil-nutrient turn-over rates (Dwyer *et al*., 2015). Each block was split into 50 × 50 cm plots and each plot was further subdivided into four 25 × 25 cm quadrats. One individual of either focal species near the center of each quadrat was assigned as the focal individual for that quadrat. Which focal species were in a given quadrat depended on the natural distribution of individuals. This experiment employed five thinning treatments at the plot level to manipulate local community compositions (individual focal individuals with no competitors, native dominated competitors, exotic dominated competitors, monocultures with only conspecific competitors, and unmanipulated plots) (Wainwright *et al*., 2019). This ensured a range of observed densities of both species and the background communities to inform model estimates of competition coefficients and intrinsic growth rates. Across both reserves we used data from 129 focal individuals in 69 plots interacting with 45 neighbouring species for *W. acuminata* and 95 focal individuals in 54 plots interacting with 40 species for *A. calendula*.

We applied our sparse modeling framework to quantify the effect of the competitive environment on fecundity in *W. acuminata* and *A. calendula* under different environmental conditions. Fecundity *F_e,i_* was measured as the number of flowers produced by each focal individual. The competitive environment was characterized as the number of individuals of each interacting species in the quadrat after the experimental treatment had been applied (*N_e, j_*). We considered two aspects of the physical environment *X_e_*: percent overhead tree canopy cover, measured at the plot scale, and soil Colwell P (mg/kg), measured at the block scale. Both environmental covariates were standardized for inclusion in the model. We ran a separate model for each focal species and environmental covariate, for a total of four model fits. To account for regional differences between the Bendering and Perenjori reserves, we incorporated a fixed effect for the two different reserves into our sparse modeling approach by allowing *λ_i_*, *λ_e,i_*, 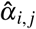, and 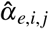 to differ between reserves. Using this approach, we quantified *λ_e,i_* and *α_e,i,j_* for both species across both environmental gradients in the York gum-jam woodland communities.

## 4 Results

### 4.1 Simulations

Our model accurately predicted growth rates for simulated communities even with relatively low sample sizes (Fig. 1) and across different model formalizations (Box 1). With only 10 full-community and 10 no-competition plots, the model predicted growth rates with a root-mean-square error (RMSE) of 0.495 (credible interval, CI: 0.353-0.665). While increasing sample size further increased model accuracy (RMSE of 0.315 (CI: 0.211-0.520) for 50 plots and 0.227 (CI 0.195-0.288) for 200 plots), these results indicate the model can accurately predict species’ realized growth rates using limited data. Furthermore, species’ growth rates can be accurately predicted using observed competitive communities paired with no-competition plots, rather than necessitating common manipulative experimental designs where each possible species combination is paired across a gradient of densities (Hallett *et al*., 2019; Kraft *et al*., 2015).

**Figure 1:**
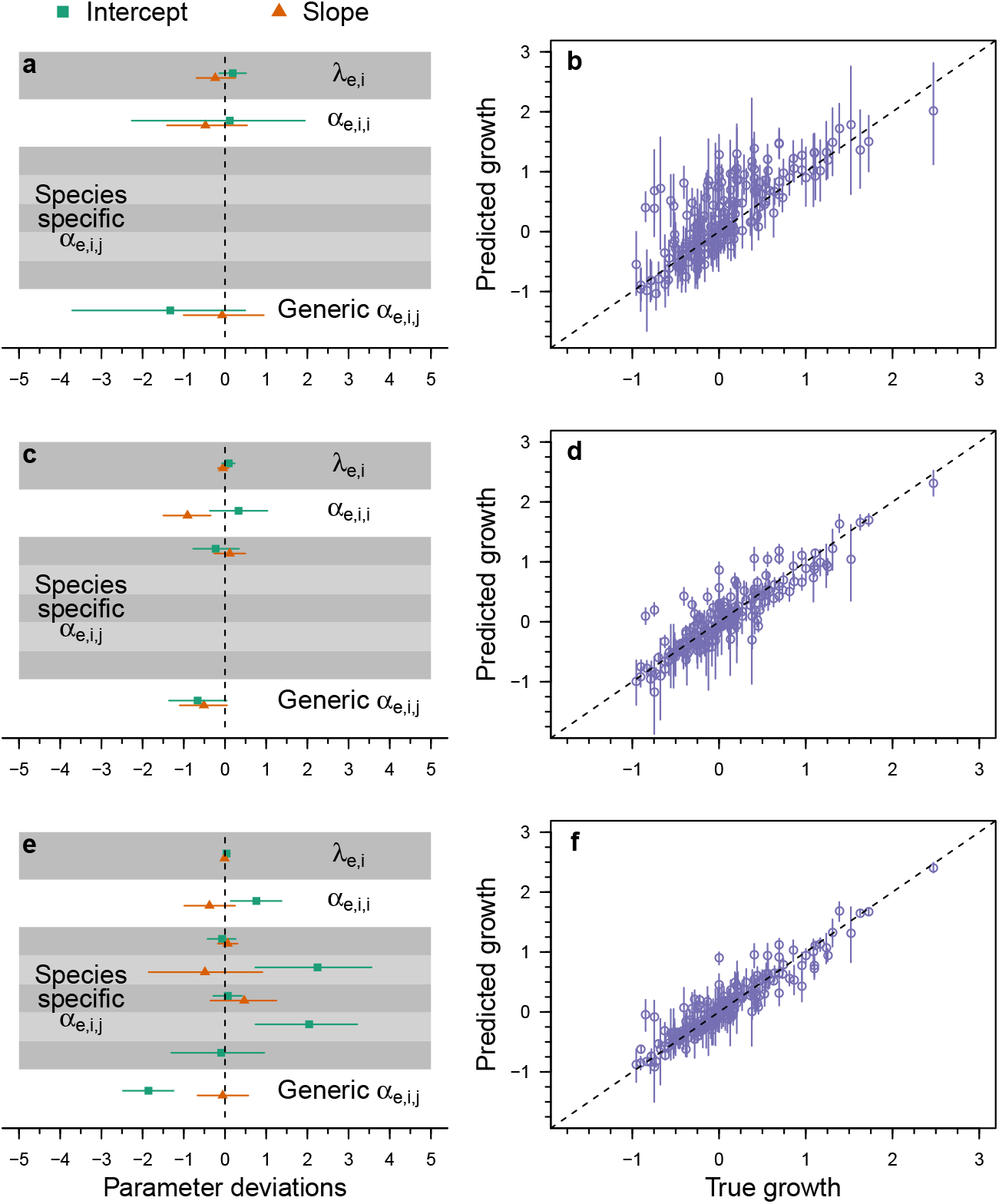
Fitted model parameter estimates and predicted growth rates. We fit the model to simulated data with 10 (a and b), 50 (c and d), and 200 (e and f) full-community and no-competition plots. The left column (a, c, and e) shows the deviation of parameter values from the true value used in the simulations (points are posterior means and lines are 95% credible intervals). The right column (b, d, and f) shows model accuracy of the focal species’ growth rate for 200 simulated full-community plots not included in the model fitting. Growth rates were calculated as 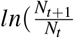. The dashed line is the 1-1 line indicating a perfect match. Points show mean estimates and lines are 95% CIs.

##### Box 1: Adapting the sparse modeling method to different ecological questions

This sparse modeling method is generalizable to a variety of underlying ecological models. The method’s flexibility allows researchers to pick and choose which parameters to include and how to specify them as best fits with their study system and questions of interest.

For example, the relationship between species’ growth rates and the environment can be modeled in multiple ways. A monotonic relationship would be appropriate for a study concentrated within a small spatial scale, while a humped-shape relationship would match expectations for a study over a broad environmental gradient. To demonstrate how our method can be modified for different underlying ecological models, we simulated environmental responses in growth rate two ways: with a monotonic relationship between species and the environmental conditions *λ_e,i_* and with a curved environmental optimum with a defined niche breadth for each species 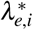. This resulted in two model formulations:

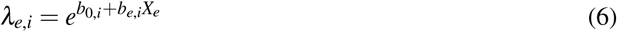

in which *b*_0,*i*_ is the mean intrinsic growth rate and *b_e,i_* is the slope of the environmental response, and

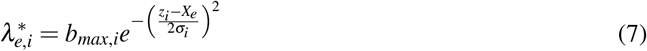

in which *b_max,i_* is the maximum intrinsic growth rate, *z_i_* is the environmental optimum, and *σ_i_* is the environmental niche breadth (following the parameterization in Thompson *et al*. (2020)). We tested these models using samples of 50 full community plots and 50 no-competition plots.

All growth rate parameters fell within the 95% credible intervals for parameters in both models. In the monotonic *λ_e,i_* model, both the intercept *b*_0,*i*_ and the slope *b_e,i_* deviated from the true values by 3%. In the optimum 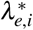 model, the maximum *b_max,i_* deviated from the true value by 6%, the niche breadth *σ_i_* deviated by 1%, and the location of the environmental optimum *z_i_* deviated by 13%, which is an absolute difference of 0.04 (Fig. 2a and b).

As a further example, the species interaction components of the model can be adjusted depending on the main research questions of interest. For questions focused on species interactions, modeling interaction coefficients independent of environmental conditions would optimize the number of non-generic species pairs identified from a given sample size of data. We compared a simple simulation with competitive interactions independent of environmental conditions *α_i,j_* to a more complex model with competitive interactions dependent on the environmental conditions *α_e,i,j_*, and modeled each accordingly. We tested these models using different sample sizes of full community data points and focal individuals. Both models predicted true out-of-sample growth rates with average RMSE of ≈ 0.4 with only 10 full-community and no-competition plots. With 80 or more full-community and no-competition plots the simpler model had a RMSE of ≈ 0.2 and the model of the more complex simulation had a RMSE of ≈ 0.25 (Figure 2c). The simpler model without species-environment interactions highlighted five non-generic *â*_0,*i*,*j*_ species pair interactions with just 50 full-community and no-competition plots. The more complex model highlighted only two non-generic terms with 50 full-community and no-competition plots. At higher sample sizes the number of non-generic terms was constrained by the global shrinkage parameter *τ* (Figure 2d)

We present these options as launching-off points for researchers to adapt the sparse modeling approach to their study systems and questions. Even more extensive modifications are possible; for example, replacing the Beverton-Holt community framework with a different underlying ecological model.

Our model was also able to accurately predict individual parameter estimates for simulated communities (Fig. 1). In particular, estimates of the intercept and slope parameters for intrinsic growth rate (*b*_0_, *i* and *b_e_*, *i* respectively) dramatically increased in accuracy from 10 to 200 data points (Fig. 1a,e). The accuracy of the estimates for the slope and intercept of intraspecific competition (*a*_0,*i*,*i*_ and *a_e,i,i_* respectively) also increased with more data, but less dramatically than the terms defining intrinsic growth rate. Parameters associated with interspecific competition (*ā*_0,*i*_ representing the intercept and *ā_e,i_* for the slope) also increased in accuracy with increasing data, although there was more variance in this relationship. This is likely because the model correctly identified a larger number of species-specific terms with more data, which decreased the total number of species contributing to the estimation of *ā*_0,*i*_ and *ā_e,i_*. When fit to only 10 simulated plots, the model did not identify any species-specific terms (*â*_0,*i*,*j*_ or *â*_*e*,*i*,*j*_) and only used *ā*_0,*i*_ and *ā_e,i_*. The model identified two species-specific terms within a single species when fit to 50 plots and eight species-specific terms across six species when fit to 200 plots. In general, the estimates of species-specific terms were highly accurate; only two out of the eight estimated species-specific interaction terms did not include the true value in their 95% credible intervals.

### 4.2 Empirical application

Our method identified environmental dependencies in intrinsic growth rates (Fig. 3a,b; 4, a,b), relative strengths of intraspecific competition and average interspecific competition, along with competition-environment interactions (Fig. 3c,d; 4, c,d), all of which differed between our two focal species. Additionally, our model highlighted three species with deviations from the average interspecific effects on native *W. acuminata*, but no such species when fit to data on exotic *A. calendula*.

*W. acuminata* and *A. calendula*’s intrinsic growth rates differed in their relationship with the environmental gradients and reserves. The intrinsic growth rate of *W. acuminata* across both environmental gradients varied between the Bendering and Perenjori reserves (Fig. 3a,b). In contrast, *λ_e,i_* for *A. calendula* was quite similar between the two reserves as it varied with both phosphorous and canopy cover (Fig. 4a,b). This could reflect local adaptation in regional populations of the native *W. acuminata* but not in the newly introduced *A. calendula*. Importantly, the intrinsic growth rate of *W. acuminata* declined with high phosphorous (marginally in Bendering, but substantially in Perenjori) while *A. calendula*’s intrinsic growth rate increased with phosphorous, potentially explaining the high prevalence of invasive species in areas with increased phosphorous (Dwyer *et al*., 2015).

Relative effects of competition between conspecifics versus heterospecifics also differed between the two focal species. For *W. acuminata*, the relationship between intraspecific competition and average interspecific competition varied with the underlying environmental gradients. At low levels of phosphorous and high levels of canopy cover, intraspecific competition in *W. acuminata* was greater than average interspecific competition (Fig. 3c,d). However, at high levels of phosphorous and low levels of canopy cover, intra- and interspecific competition converged to similar values. On the other hand, intraspecific competition for *A. calendula* was similar to or lower than generic interspecific competition across both environmental gradients (Fig. 4c,d). This likely contributes to the invasive status of *A. calendula* in this ecosystem, whereas *W. acuminata* populations self-regulate under certain environmental conditions—a necessary component of stable coexistence.

Our model highlighted multiple species with competitive effects on *W. acuminata* that differed from the generic interaction term. Across the observed gradient in phosphorous, *Hyalosperma glutinosum* had a higher than average effect on *W. acuminata* in the Perenjori reserve while *Schoenus nanus* had a lower than average effect in the Bendering reserve (Fig. 3c). Across the observed gradient in canopy cover, *Hypochaeris glabra* had a much higher than average effect on *W. acuminata* in Bendering (Fig. 3d). In contrast, all heterospecific interactive effects on *A. calendula* remained grouped in the generic competition term. The lack of species with unique effects on *A. calendula* (Fig. 4c,d) could be due to its exotic status (Lai *et al*., 2015). With no shared evolutionary history with any other community members, *A. calendula* could be experiencing a form of competitive release, wherein the identity of competitor species matters less than simply the presence of additional individuals.

## 5 Discussion

Given the inherent complexity of ecological communities, ecologists are often forced to rely on simplifying assumptions in order to perform tractable analyses, such as limiting the number of species considered or ignoring environmental variation. The sparse modeling approach presented here provides an alternative method to analyze community data without requiring extensive additional data or sacrificing complexity. This approach enabled us to accurately predict population growth rates with limited data and identify how species’ demographic rates and competitive interactions depend on the environment. Our results identify environment by species interactions that deviate from the species-averaged community effects without making *a priori* assumptions about species groupings (Figure 3c, d). This information and output from the sparse modeling approach generates concrete, testable hypotheses about species interactions and environmental conditions. We see broad potential for this method’s implementation in community ecology, from theory development to management applications.

The sparse modeling approach’s flexibility in modeling populations and communities allows easy adjustments for the best match between underlying model structure and the given study system and research questions. As we show in Box 1 and Fig. 2, these models can successfully be applied to different forms of species-environment interactions and be modified to be more or less complex based on underlying ecological questions and data availability. For example, the functional form of the relationship between intrinsic growth rate and the environment likely depends on a study’s spatial scale. For localized studies, a simple monotonic relationship (Figure 2a) might be appropriate to capture species’ expected responses across a small range of environmental variation. However, studies over larger spatial scales might require a functional form with optimal intrinsic growth reached at an intermediate environmental value and declining away from that value (Figure 2b), mimicking expected patterns of adaptation across species’ ranges (Angert *et al*., 2020). Additionally, while we used a Beverton-Holt framework in our examples (Beverton & Holt, 1957), the sparse approach is agnostic to the underlying ecological model. Thus, it could be used with different functional forms of competition (García-Callejas *et al*., 2020) with models incorporating both competitive and facilitative interactions (Stachowicz, 2001) or different underlying demography such as seed banks.

**Figure 2:**
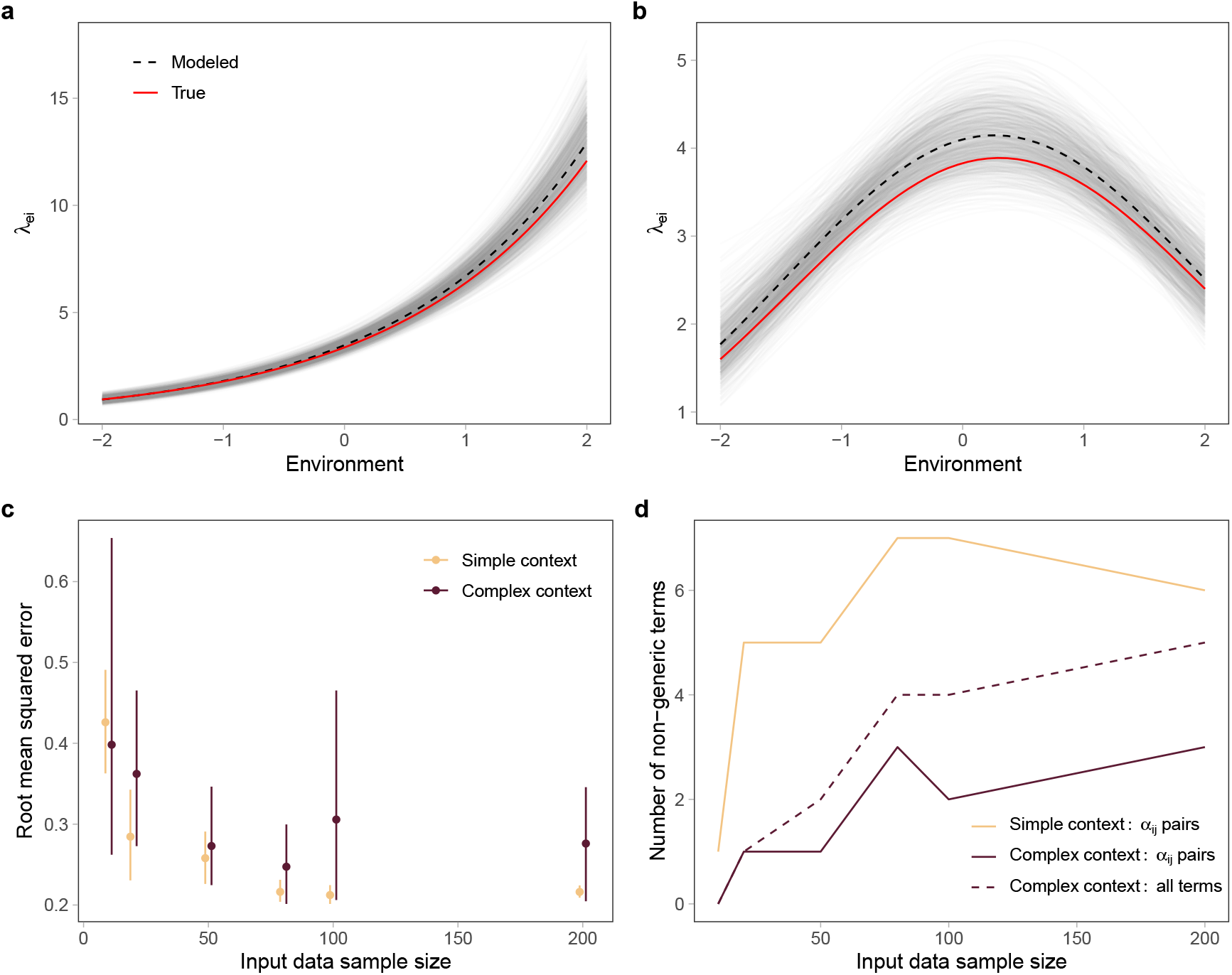
Estimates for *λ_e,i_* for simulated data with (A) a monotonic response to the environment and (B) an environmental optimum, with the true values as a solid red line, model means as a dashed black line, and individual model posterior draws as thin grey lines. Both models were run with 50 full community plots and 50 no-competition plots. All growth rate parameters fell within the 95% credible intervals for parameters in both models. (C) Simulated contexts with only *α_i,j_* species pair intercepts compared to contexts with *α_e,i,j_* species pair by environment slopes. Performance of sparse models matching these simulation contexts was measured as RMSE between the model predictions and the true value of 200 out-of-sample data points. Both models predict true growth rates similarly well at low sample sizes (10 samples) but the model of the simpler simulation converges closer to the true values. (D) Number of non-generic terms identified by models on the different *α* contexts, which is constrained at larger sample sizes by the global shrinkage parameter *τ*. The model of the species pair intercept only simulation identifies more non-generic *α_i,j_* pairs at all sample sizes. The model of the simulation with both non-generic species pair *α*_0,*i*,*j*_ intercepts and species pair by environment *α_e,i,j_* slopes requires more data to identify non-generic terms, and those are split between the intercept and slope terms.

With this flexibility, sparse modeling has the potential to be a powerful tool to accelerate the development of community ecology theory and practice. It can provide important insights into the covariation of environmental conditions, species’ demographic rates, and competitive effects—critical aspects of modern coexistence theory (Chesson, 2000). This includes quantifying the relative strengths of intra-versus inter-specific competition, which is a key condition for stable coexistence (Adler *et al*., 2018; Chesson, 2000). Furthermore, the approach elucidates the effect of environmental conditions on species’ density-independent growth rates versus competitive interactions, potentially allowing for quantification of variation-dependent coexistence mechanisms, such as the storage effect, in diverse communities (Chesson, 2000). Similarly, output from our sparse modeling approach across environmental gradients can be used to quantify the relative importance of environmental (abiotic) filtering, biotic interactions, and the joint effect on species occurrence (Cadotte & Tucker, 2017). Applying such an approach is especially exciting for linking community theory to global change predictions, depending on the underlying environmental gradient of interest.

In addition to expanding theory, we see exciting potential for sparse modeling to address questions in applied contexts and generate new hypotheses from existing datasets that inform management strategies. This includes quantifying how environmental modifications can be used in conjunction with community manipulations to control invasive species or promote native species. For example, our results from the York gum-jam woodlands of Western Australia suggest the native *W. acuminata* experiences declining fitness with increasing levels of phosphorous, particularly in the Perenjori reserve (Fig. 3a). At the same time, the model identified *H. glutinosum* as having a stronger than average competitive impact on *W. acuminata* in Perenjori (Fig. 3c). Taken together, these results suggest that reserve managers could help maintain or expand populations of *W. acuminata* by mitigating phosphorous run-off while simultaneously removing *H. glutinosum* in key locals. In contrast, our results for the invasive *A. calendula* suggest that neighbor species identity is unimportant (Fig. 4c,d) and management strategies focusing solely on environmental factors would be most impactful.

**Figure 3:**
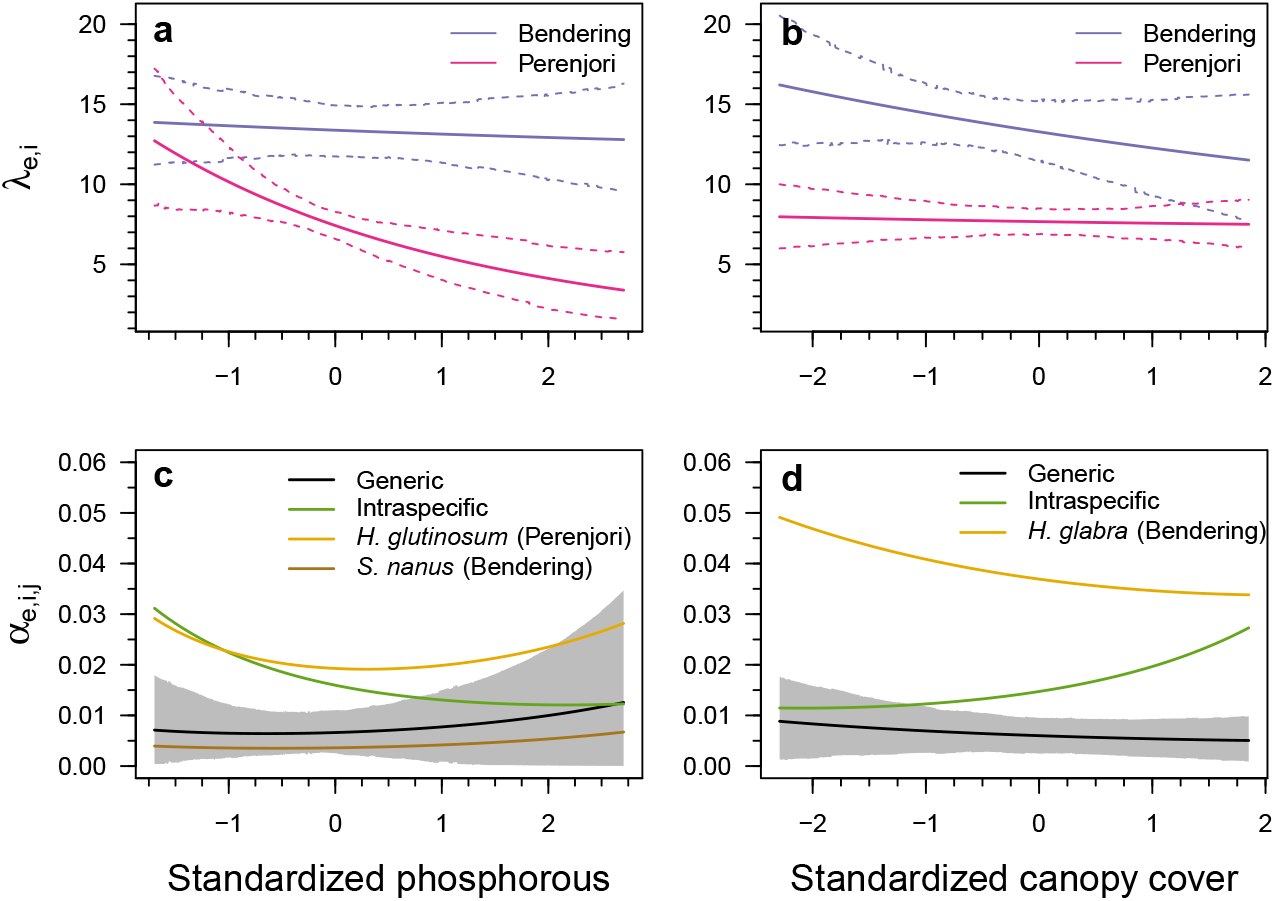
Model estimates for *W. acuminata*. Means (solid lines) and 95% CIs (dashed lines) are shown for *λ_e,i_* across a gradient of phosphorous (a) and canopy cover (b). Colors indicate the Bendering and Perenjori reserves. The mean (black line) and 95% CI for generic interspecific competition are shown across a phosphorous (c) and canopy cover (d) gradient. In both c and d, the mean intraspecific competition coefficient is shown in green and different species identified by the model as non-generic in each reserve are shown with other colors as indicated in the legends.

**Figure 4:**
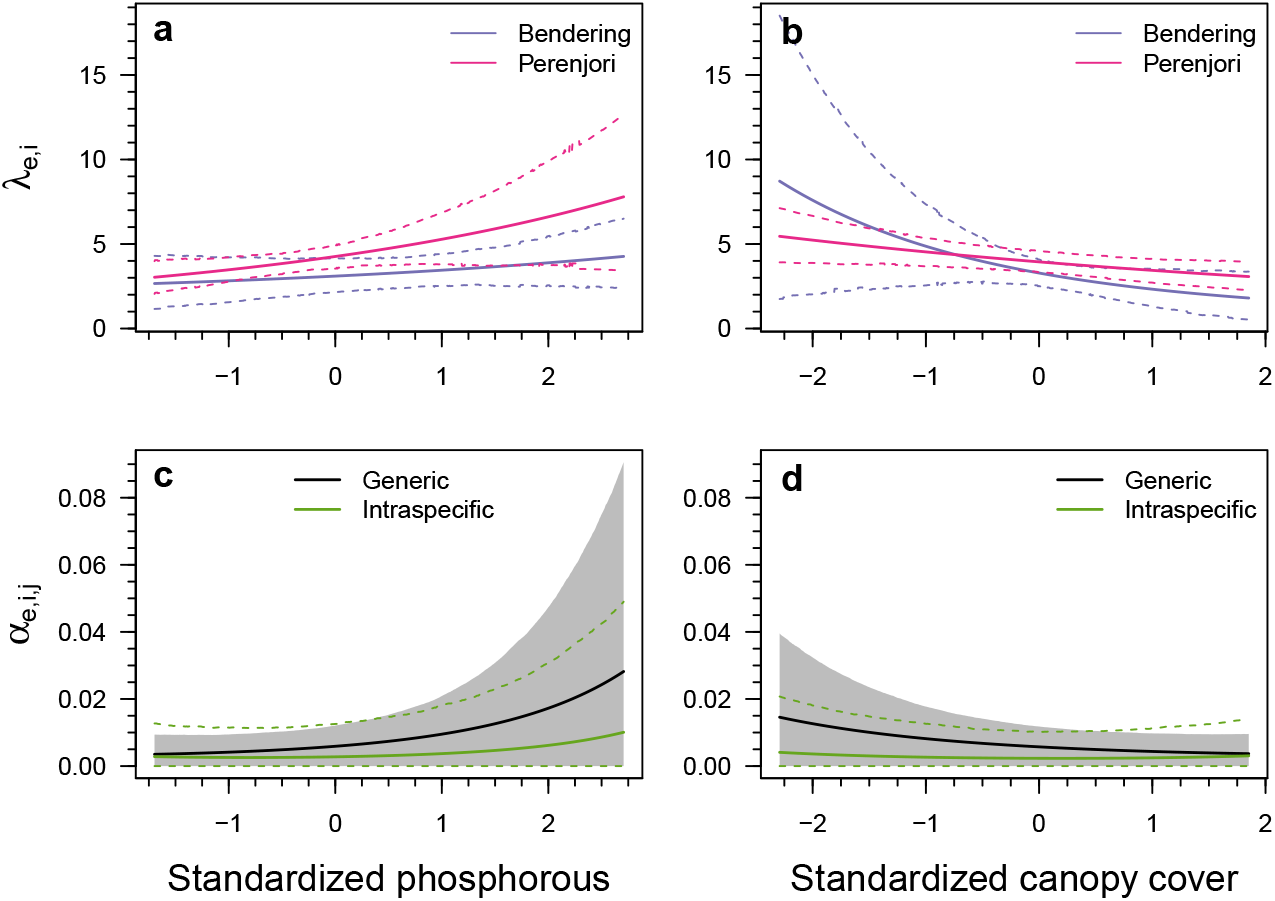
Model estimates for *A. calendula*. Means (solid lines) and 95% CIs (dashed lines) are shown for *λ_e,i_* across a gradient of phosphorous (a) and canopy cover (b). Colors indicate the Bendering and Perenjori reserves. The mean (black line) and 95% CI for generic interspecific competition are shown across a phosphorous (c) and canopy cover (d) gradient. In both c and d, the mean intraspecific competition coefficient is shown in green with dashed lines indicating the CI. The model did not identify any interspecific competitors in either reserve that impacted *A. calendula* differently from a generic competition coefficient.

Beyond the implementation of the sparse modeling approach presented here, the underlying model structure can be further adjusted to align the model with focal management questions. For example, if a management goal only requires knowledge of species interactions within a community, the model could be simplified to remove environmental covariates (Box 1). Alternatively, the global shrinkage parameter *τ* could be set at a fixed value to induce more or less sparsity in the final model results. Such a change could allow users to manually explore the trade-off between inclusion of species-specific terms and precision of parameter estimates, finding the balance that best suits their particular goals. For example, fixing *τ* to a higher value would yield more estimates of species-specific parameters, which could help to inform future research priorities, but those estimates would likely be less precise, limiting their utility in predicting community dynamics. Small adjustments such as these empower ecologists and managers to match the tool to their questions and aims.

In its current structure, the sparse modeling framework is most useful when applied to high-diversity communities with limited available data. As we observed when analyzing simulated data, the number of non-generic terms does not necessarily increase with sample size, and is limited by *τ* at higher sample sizes (Figure 2d). As described above, there may be cases where manual adjustments to *τ* would be beneficial depending on the available data and questions of interest. However, a traditional, non-sparse model in which every interaction term is included may still be preferable in situations with abundant data, lowerdiversity communities, or when answering questions requiring individual estimates of all potential species interactions. In contrast, the sparse approach is particularly helpful with limited data and in cases where traditional models often struggle to converge or provide overly broad parameter estimates.

The model we present here is currently analyzed for a single growing season and for use with traditional population dynamic models (e.g. Beverton-Holt or Lotka-Voltera models). Given the importance of temporal stochasticity to community dynamics (Shoemaker *et al*., 2020) and the need to predict community responses to changing anthropogenic pressures (Ma *et al*., 2017), sparse modeling with time series could provide invaluable insight into the importance of species-specific interactions through time as well as space. Further, extending the approach to a wider range of input data beyond individual counts (e.g. percent cover or biomass) would allow for future uses across observational datasets and especially for perennial-dominated systems.

Sparse modeling approaches have proved immensely valuable in fields as diverse as genomics (Gianola & Fernando, 2020) to economics (Fan *et al*., 2011). By dramatically reducing the parameter load required to model diverse communities across environmental gradients, we show these sparse modeling approaches can provide both theoretical and applied insights in community ecology as well. We demonstrate the flexibility of this approach across different ecological models and underlying biological assumptions, and are excited to see it expanded and applied to a variety of ecological questions and applications. Although the implementation of the sparse method requires an initial conceptual investment, the output results are easily interpretable—a quality that is particularly important for linking models to practice. The sparse modeling approach eliminates the need for *a priori* assumptions regarding species’ groupings or the exclusion of all but a handful of focal species, providing a critical method and step forward in expanding ecological theory and linking models to observational and experimental datasets of diverse communities.

## 6 Acknowledgements

This paper is a joint effort of the working group sToration kindly supported by sDiv, the Synthesis Centre of the German Centre for Integrative Biodiversity Research (iDiv) Halle-Jena-Leipzig, funded by the German Research Foundation (FZT 118, 02548816). CWL, CMW, LGS were supported by Modelscapes, NSF award #EPS-2019528. OG was supported by the Spanish Ministry of Economy and Competitiveness (MINECO) and by the European Social Fund through the Ramón y Cajal Program (RYC-2017-23666); GB was supported by the Swedish Research Council (Vetenskapsra°det), grant 2017-05245. MMM was supported by the Australian Research Council (DP140100574).

## Literature Cited

Adler, P. B., Smull, D., Beard, K. H., Choi, R. T., Furniss, T., Kulmatiski, A., Meiners, J. M., Tredennick, A. T. & Veblen, K. E. (2018). Competition and coexistence in plant communities: intraspecific competition is stronger than interspecific competition. Ecology letters, 21, 1319–1329.

Allesina, S. & Levine, J. M. (2011). A competitive network theory of species diversity. Proceedings of the National Academy of Sciences, 108, 5638–5642.

Angert, A. L., Bontrager, M. G. & Ågren, J. (2020). What do we really know about adaptation at range edges? Annual Review of Ecology, Evolution, and Systematics, 51, 341–361.

Barner, A. K., Coblentz, K. E., Hacker, S. D. & Menge, B. A. (2018). Fundamental contradictions among observational and experimental estimates of non-trophic species interactions. Ecology, 99, 557–566.

Beverton, R. J. & Holt, S. J. (1957). On the dynamics of exploited fish populations, vol. 11. Ministry of Agriculture, Fisheries and Food.

Bhadra, A., Datta, J., Polson, N. G., Willard, B. et al. (2019). Lasso meets horseshoe: A survey. Statistical Science, 34, 405–427.

Bimler, M. D., Stouffer, D. B., Lai, H. R. & Mayfield, M. M. (2018). Accurate predictions of coexistence in natural systems require the inclusion of facilitative interactions and environmental dependency. Journal of Ecology, 106, 1839–1852.

Bulleri, F., Bruno, J. F., Silliman, B. R. & Stachowicz, J. J. (2016). Facilitation and the niche: implications for coexistence, range shifts and ecosystem functioning. Functional Ecology, 30, 70–78.

Cadotte, M. W. & Tucker, C. M. (2017). Should environmental filtering be abandoned? Trends in ecology & evolution, 32, 429–437.

Carvalho, C. M., Polson, N. G. & Scott, J. G. (2009). Handling sparsity via the horseshoe. In: Artificial Intelligence and Statistics.

Chesson, P. (2000). Mechanisms of maintenance of species diversity. Annual Review of Ecology and Systematics, 31, 343–366.

Clark, A. T., Ann Turnbull, L., Tredennick, A., Allan, E., Harpole, W. S., Mayfield, M. M., Soliveres, S., Barry, K., Eisenhauer, N., de Kroon, H. et al. (2020a). Predicting species abundances in a grassland biodiversity experiment: Trade-offs between model complexity and generality. Journal of ecology, 108, 774–787.

Clark, J. S., Scher, C. L. & Swift, M. (2020b). The emergent interactions that govern biodiversity change. Proceedings of the National Academy of Sciences, 117, 17074–17083.

Dwyer, J. M., Hobbs, R. J., Wainwright, C. E. & Mayfield, M. M. (2015). Climate moderates release from nutrient limitation in natural annual plant communities. Global Ecology and Biogeography, 24, 549–561.

Fan, J., Lv, J. & Qi, L. (2011). Sparse high-dimensional models in economics. Annu. Rev. Econ., 3, 291–317.

García-Callejas, D., Godoy, O. & Bartomeus, I. (2020). cxr: A toolbox for modelling species coexistence in r. Methods in Ecology and Evolution, 11, 1221–1226.

Germain, R. M., Mayfield, M. M. & Gilbert, B. (2018). The ‘filtering’metaphor revisited: competition and environment jointly structure invasibility and coexistence. Biology letters, 14, 20180460.

Gianola, D. & Fernando, R. L. (2020). A multiple-trait bayesian lasso for genome-enabled analysis and prediction of complex traits. Genetics, 214, 305–331.

Godoy, O. & Levine, J. M. (2014). Phenology effects on invasion success: insights from coupling field experiments to coexistence theory. Ecology, 95, 726–736.

Hallett, L. M., Shoemaker, L. G., White, C. T. & Suding, K. N. (2019). Rainfall variability maintains grass-forb species coexistence. Ecology Letters, 22, 1658–1667.

Hastie, T., Tibshirani, R. & Wainwright, M. (2015). Statistical learning with sparsity: the lasso and generalizations. CRC press.

HilleRisLambers, J., Adler, P. B., Harpole, W., Levine, J. M. & Mayfield, M. M. (2012). Rethinking community assembly through the lens of coexistence theory. Annual review of ecology, evolution, and systematics, 43, 227–248.

König, C., Wüest, R. O., Graham, C. H., Karger, D. N., Sattler, T., Zimmermann, N. E. & Zurell, D. (2021). Scale dependency of joint species distribution models challenges interpretation of biotic interactions. Journal of Biogeography.

Kraft, N. J., Godoy, O. & Levine, J. M. (2015). Plant functional traits and the multidimensional nature of species coexistence. Proceedings of the National Academy of Sciences, 112, 797–802.

Kühner, A. & Kleyer, M. (2008). A parsimonious combination of functional traits predicting plant response to disturbance and soil fertility. Journal of Vegetation Science, 19, 681–692.

Lai, H. R., Mayfield, M. M., Gay-des combes, J. M., Spiegelberger, T. & Dwyer, J. M. (2015). Distinct invasion strategies operating within a natural annual plant system. Ecology Letters, 18, 336–346.

Lanuza, J. B., Bartomeus, I. & Godoy, O. (2018). Opposing effects of floral visitors and soil conditions on the determinants of competitive outcomes maintain species diversity in heterogeneous landscapes. Ecology Letters, 21, 865–874.

Legendre, P. & Gauthier, O. (2014). Statistical methods for temporal and space–time analysis of community composition data. Proceedings of the Royal Society B: Biological Sciences, 281, 20132728.

Letten, A. D., Dhami, M. K., Ke, P.-J. & Fukami, T. (2018). Species coexistence through simultaneous fluctuation-dependent mechanisms. Proceedings of the National Academy of Sciences, 115, 6745–6750.

Levine, J. M. & HilleRisLambers, J. (2009). The importance of niches for the maintenance of species diversity. Nature, 461, 254.

Levins, R. & Culver, D. (1971). Regional coexistence of species and competition between rare species. Proceedings of the National Academy of Sciences, 68, 1246–1248.

Li, Y., Mayfield, M. M., Wang, B., Xiao, J., Kral, K., Janik, D., Holik, J. & Chu, C. (2021). Beyond direct neighbourhood effects: higher-order interactions improve modelling and predicting tree survival and growth. National Science Review, 8, nwaa244.

Ma, Z., Liu, H., Mi, Z., Zhang, Z., Wang, Y., Xu, W., Jiang, L. & He, J.-S. (2017). Climate warming reduces the temporal stability of plant community biomass production. Nature Communications, 8, 1–7.

Martyn, T. E., Stouffer, D. B., Godoy, O., Bartomeus, I., Pastore, A. & Mayfield, M. M. (2020). Identifying ‘useful’fitness models: balancing the benefits of added complexity with realistic data requirements in models of individual plant fitness. the American Naturalist.

May, R. M. & Leonard, W. J. (1975). Nonlinear aspects of competition between three species. SIAM journal on applied mathematics, 29, 243–253.

Mayfield, M. M. & Levine, J. M. (2010). Opposing effects of competitive exclusion on the phylogenetic structure of communities. Ecology letters, 13, 1085–1093.

Mayfield, M. M. & Stouffer, D. B. (2017). Higher-order interactions capture unexplained complexity in diverse communities. Nature ecology & evolution, 1, 1–7.

O’Hara, R. B., Sillanpää, M. J. et al. (2009). A review of bayesian variable selection methods: what, how and which. Bayesian analysis, 4, 85–117.

Ovaskainen, O., Rybicki, J. & Abrego, N. (2019). What can observational data reveal about metacommunity processes? Ecography, 42, 1877–1886.

Ovaskainen, O., Tikhonov, G., Dunson, D., Grøtan, V., Engen, S., Sæther, B.-E. & Abrego, N. (2017a). How are species interactions structured in species-rich communities? a new method for analysing time-series data. Proceedings of the Royal Society B: Biological Sciences, 284, 20170768.

Ovaskainen, O., Tikhonov, G., Norberg, A., Guillaume Blanchet, F., Duan, L., Dunson, D., Roslin, T. & Abrego, N. (2017b). How to make more out of community data? a conceptual framework and its implementation as models and software. Ecology letters, 20, 561–576.

Pérez-Ramos, I. M., Matías, L., Gómez-Aparicio, L. & Godoy, Ó. (2019). Functional traits and phenotypic plasticity modulate species coexistence across contrasting climatic conditions. Nature communications, 10, 1–11.

Picoche, C. & Barraquand, F. (2020). Strong self-regulation and widespread facilitative interactions in phytoplankton communities. Journal of Ecology, 108, 2232–2242.

Piironen, J., Vehtari, A. et al. (2017). Sparsity information and regularization in the horseshoe and other shrinkage priors. Electronic Journal of Statistics, 11, 5018–5051.

R Core Team (2019). R: A Language and Environment for Statistical Computing. R Foundation for Statistical Computing, Vienna, Austria. URL https://www.R-project.org/.

Ricker, W. E. (1954). Stock and recruitment. Journal of the Fisheries Board of Canada, 11, 559–623.

Schamp, B. S., Arnott, S. E. & Joslin, K. L. (2015). Dispersal strength influences zooplankton co-occurrence patterns in experimental mesocosms. Ecology, 96, 1074–1083.

Shoemaker, L. G. & Melbourne, B. A. (2016). Linking metacommunity paradigms to spatial coexistence mechanisms. Ecology, 97, 2436–2446.

Shoemaker, L. G., Sullivan, L. L., Donohue, I., Cabral, J. S., Williams, R. J., Mayfield, M. M., Chase, J. M., Chu, C., Harpole, W. S., Huth, A. et al. (2020). Integrating the underlying structure of stochasticity into community ecology. Ecology, 101, e02922.

Spaak, J. W. & De Laender, F. (2020). Intuitive and broadly applicable definitions of niche and fitness differences. Ecology letters, 23, 1117–1128.

Stachowicz, J. J. (2001). Mutualism, facilitation, and the structure of ecological communities: positive interactions play a critical, but underappreciated, role in ecological communities by reducing physical or biotic stresses in existing habitats and by creating new habitats on which many species depend. Bioscience, 51, 235–246.

Stan Development Team (2018). RStan: the R interface to Stan. URL http://mc-stan.org/. R package version 2.18.2.

Suppiah, R., Hennessy, K., Whetton, P., McInnes, K., Macadam, I., Bathols, J., Ricketts, J. & Page, C. (2007). Australian climate change projections derived from simulations performed for the ipcc 4th assessment report. Australian Meteorological Magazine, 56, 131–152.

Thompson, P. L., Guzman, L. M., De Meester, L., Horváth, Z., Ptacnik, R., Vanschoenwinkel, B., Viana, D. S. & Chase, J. M. (2020). A process-based metacommunity framework linking local and regional scale community ecology. Ecology letters, 23, 1314–1329.

Tilman, D. (1982). Resource competition and community structure. Princeton university press.

Tredennick, A. T., Hooker, G., Ellner, S. P. & Adler, P. B. (2021). A practical guide to selecting models for exploration, inference, and prediction in ecology. Ecology, e03336.

Uriarte, M., Condit, R., Canham, C. D. & Hubbell, S. P. (2004). A spatially explicit model of sapling growth in a tropical forest: does the identity of neighbours matter? Journal of Ecology, 92, 348–360.

Van Erp, S., Oberski, D. L. & Mulder, J. (2019). Shrinkage priors for bayesian penalized regression. Journal of Mathematical Psychology, 89, 31–50.

Vellend, M. (2020). The theory of ecological communities (MPB-57), vol. 57. Princeton University Press.

Wainwright, C. E., HilleRisLambers, J., Lai, H. R., Loy, X. & Mayfield, M. M. (2019). Distinct responses of niche and fitness differences to water availability underlie variable coexistence outcomes in semi-arid annual plant communities. Journal of Ecology, 107, 293–306.

